# Molecular and cellular phenotypic differences distinguish murine syngeneic models from human tumors

**DOI:** 10.1101/694232

**Authors:** Wenyan Zhong, Jeremy S. Myers, Fang Wang, Kai Wang, Justin Lucas, Edward Rosfjord, Judy Lucas, Andrea T. Hooper, Sharon Yang, LuAnna Lemon, Magali Guffroy, Chad May, Jadwiga R. Bienkowska, Paul A. Rejto

## Abstract

The clinical success of immune checkpoint inhibitors that target cytotoxic T-lymphocyte associated protein 4 (CTLA4) and programmed cell death protein 1 (PD-1) or programmed death ligand-1 (PD-L1) demonstrates that reactivation of the human immune system delivers durable responses for some patients and represents an exciting approach for cancer treatment. The combination of multiple immunotherapies as well as the combination of immunotherapy with targeted therapy is being pursued vigorously to increase the rate and extend the duration of response. Preclinical *in vivo* models for immuno-oncology (IO) typically require immunocompetent mice bearing murine syngeneic tumors. To facilitate translation of preclinical studies into human, we characterized the genomic, transcriptomic, and protein expression of a panel of mouse tumor cell lines grown *in vitro* culture as well as *in vivo* tumor samples. Our studies identified many genetic and cellular phenotypic differences that distinguish murine syngeneic models from human cancers. For example, only a small fraction of the somatic single nucleotide variants (SNVs) in mouse cell lines directly match SNVs from human actionable cancer genes. At the cellular level, some epithelial tumor models have a more mesenchymal phenotype with relatively low T-lymphocyte infiltration compared to the corresponding human cancers. Furthermore, in contrast to what has been reported for human tumors, we did not observe a correlation between neoantigen load and cytolytic activity in syngeneic models. Finally, the relative immunogenicity of syngeneic tumors does not typically resemble that of human tumors of the same tissue origin. CT26, a colon tumor model, had the highest immunogenicity and was the most responsive model to CTLA4 inhibitor treatment, by contrast to the relatively low immunogenicity and response rate to checkpoint inhibitor therapies in human colon cancers. These differences highlight limitations of syngeneic models for evaluating novel immune therapies and rationalize some of the challenges associated with translating preclinical findings to clinical studies.

## Introduction

Preclinical mouse models support cancer therapeutic development by contributing to target validation, elucidation of drug mechanism of action, and generation of biomarker hypotheses to test in clinical settings. Two major categories of preclinical mouse models are immune compromised and immune competent (Gould et al. 2015). Patient-derived xenografts (PDXs) and cell-line derived xenografts (CDXs) originate by transplanting either human tumor explants or established human tumor cell lines into immune deficient mouse hosts. These models have been widely applied in developing cancer therapies that modulate tumor cell autonomous functions. The rich genetic information about cancer cell lines (Barretina et al. 2012) and PDXs (Gao et al. 2015; Gu et al. 2015) from extensive genomic characterization supports model selection to investigate specific target biology or perform drug sensitivity screens. Unfortunately CDXs and PDXs have limited use in cancer immunotherapy evaluation because an immune compromised host is required for xenotransplantation. By contrast, immune competent mouse model systems such as syngeneic mouse models, derived by transplanting established mouse cell lines or tumor tissues to strain-matched mouse hosts, and genetically engineered mouse models (GEMMs), created by introducing genetic modifications that result in spontaneous tumor development, retain intact murine immune systems and are better suited to study the interplay between immune and tumor cells.

The recent approval of immune checkpoint inhibitors and their success in generating durable response in some patients has reinvigorated interest in developing novel immune therapies and evaluating combination regimens (Sharma and Allison 2015). While syngeneic mouse models and GEMMs both possess intact immune systems (Dranoff 2012), GEMMs typically have few mutations and lower immunogenicity. Syngeneic mouse models, which have served as workhorses for investigating immune therapies and studying the intricate immune surveillance of cancer development, have a wide spectrum of mutations (Ostrand-Rosenberg 2004; Dranoff 2012). Anti-tumor activity via checkpoint blockade, such as with a CTLA4 blocking antibody, was initially observed in syngeneic models (Grosso and Jure-Kunkel 2013), suggesting that syngeneic model findings may translate to the clinic. The anti-CTLA4 antibody has variable response among different syngeneic models with marked response in CT26, GL261, and EMT6 models, while it was shown to be ineffective in B16F10, a melanoma model (Grosso and Jure-Kunkel 2013). This response pattern was postulated to be due to the diverse immunogenicity among the models, although the underlying molecular mechanisms remain elusive due in part to a lack of understanding of the immunogenic state that favors response. Compared to patient-derived xenograft models, there have been far fewer syngeneic models established and characterized. Recently, several studies have been published that begin to profile the molecular and cellular characteristics of murine immune competent mouse models (Lechner et al. 2013; Castle et al. 2014a; Castle et al. 2014b; Mosely et al. 2017; Yang et al. 2017).

We compared genomic, proteomic and immunohistochemistry features of a panel of ten mouse syngeneic models with human tumors in The Cancer Genome Atlas (http://cancergenome.nih.gov/). We characterized the mutational landscape and predicted the neoantigen burden of these models through whole exome sequencing, and compared the variants identified in syngeneic models to mutations in human tumors. We also evaluated syngeneic model tumor phenotypes through immunohistochemistry and compared the architecture to human cancers. We performed RNA-Seq of tumors grown in syngeneic mice as well as the same cells grown in culture, and predicted immune infiltration through computational deconvolution of gene expression data into immune components. Compared to previous studies (Lechner et al. 2013; Castle et al. 2014a; Castle et al. 2014b; Mosely et al. 2017; Yang et al. 2017), our study includes expression analysis for syngeneic models from both cells grown *in vitro* culture as well as *in vivo* tumor samples, enabling an assessment of tumor cell intrinsic properties. We also characterized these mouse syngeneic models by proteomics which enabled us to verify gene expression findings identified from transcription profiling, as well as to identify potential mouse virus proteins that may contribute to immunogenicity.

## Results

### Murine syngeneic models do not fully recapitulate mutations observed in human tumors

We performed whole exome sequencing (WES) of ten syngeneic models (Table 1, Supplemental Table S1) to assess their mutational landscape. Similar to human tumors, the majority of models have frequent missense mutations (Supplemental Figure 1). To assess the accuracy of our variant calls, we independently tested 115 variants mapped to the TARGET (tumor <u>a</u>lterations relevant for <u>ge</u>nomics-driven therapy) database (http://archive.broadinstitute.org/cancer/cga/target) by Sanger sequencing: all 115 of the predicted variants were validated (Supplemental Table S2), supporting a high level of precision for our variant calls. The transition/transversion (Ts/Tv) ratio varied across a wide span ranging from 0.27 to 3.65 (Supplemental Figure 2A), similar to the Ts/Tv range in somatic variants from human cancers (Supplemental Figure 2A) (Alexandrov et al. 2013). MC38 has many more transversion SNVs than transition SNVs, while more than 50% of the SNV for the CT26 model are C>T;G>A transition SNVs (Supplemental Figure 2B). The broad Ts/Tv range in syngeneic models may be attributed to the variety of mutagens used to derive these models and different DNA repair mechanisms. For example, MC38, a model induced by the DNA methylating agent DMH, is enriched with C>A;G>T transversion SNVs while the CT26 model was generated by the carcinogen NMU, known to induce C>T;G>A mutations (Burns et al. 1988). Previously, both transversion and transition mutations were reported to be induced by DMH in murine Trp53 genes (Jenkins et al. 1997).

**Figure 1.**
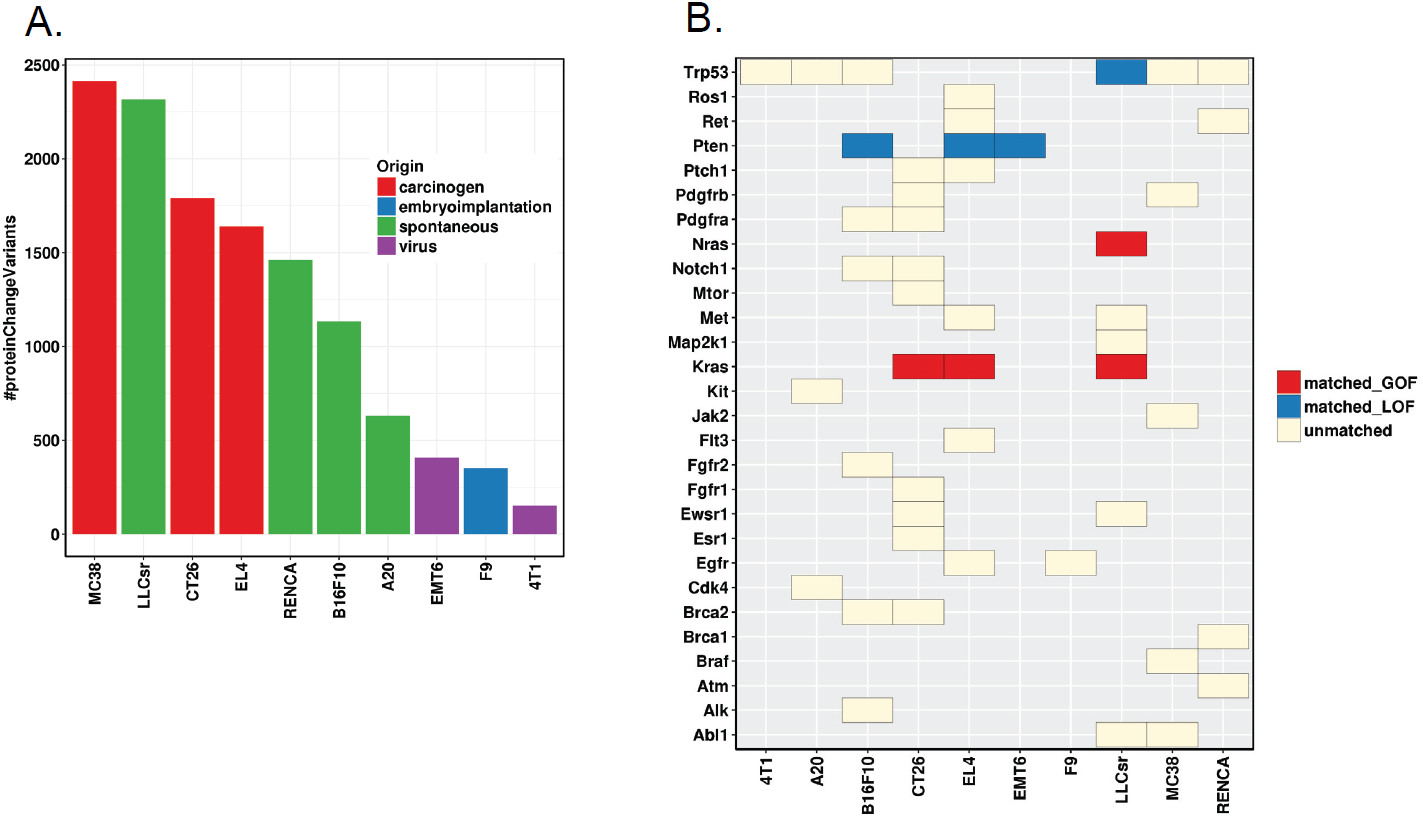
A. Variants predicted to alter protein functions (variant effect defined as MODERATE, “A non-disruptive variant that might change protein effectiveness”, or HIGH, “The variant is assumed to have disruptive impact in the protein, probably causing protein truncation, loss of function or triggering nonsense mediated decay”, by SnpEff). B. Protein sequence altering variants of known cancer genes; GOF: gain of function; LOF: loss of function; matched_GOF: mouse variants matching human GOF variants (exact variants); matched_LOF: mouse variants matching human LOF variants (truncating mutation or missense mutation at the same amino acid); unmatched: mouse variants not reported as known actionable variants in human tumors.

**Figure 2.**
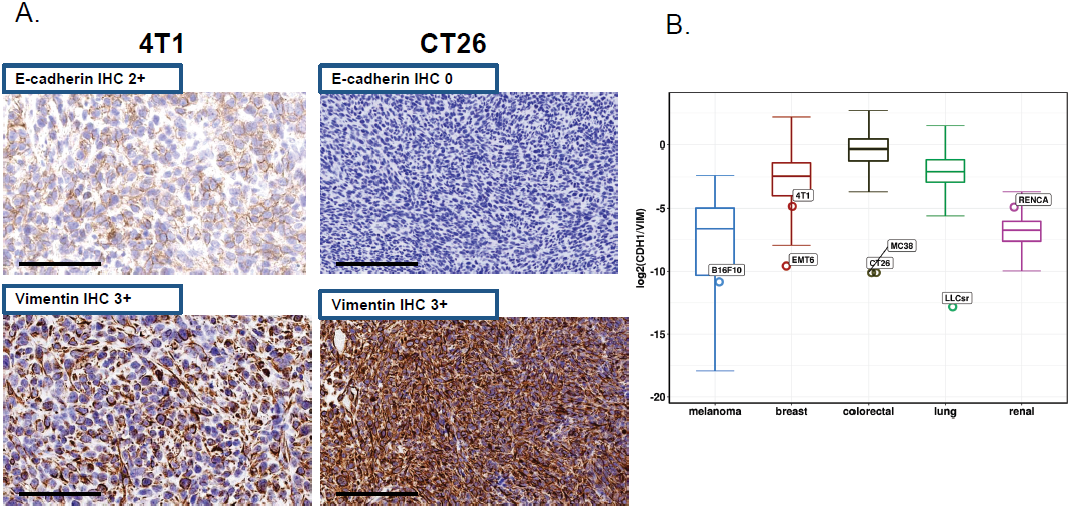
Mesenchymal-like phenotype of some syngeneic tumors. A. E-cadherin and vimentin stain in 4T1 and CT26 models. B. Comparison of Ratio of E-cadherin to vimentin between solid tumor syngeneic models (open circle) with tissue matched human tumors from TCGA (boxplot; lung: lung adenocarcinoma and lung squamous cell carcinoma). Ratio was calculated with the expression value (TPM) of E-cadherin and vimentin.

**Table 1.** Mutational load of the 10 syngeneic murine models and the corresponding human cancer (LUAD: Lung adenocarcinoma, LUSC: Lung squamous cell carcinoma).

| **Model** | **Tumor Type** | **Parent Strain** | **Origin** | **Mutational Load (per Mb)** | **Median Human Mutational Load (per Mb)** |
| --- | --- | --- | --- | --- | --- |
| **4T1** | **breast** | **BALB/c** | **virus** | **5** | **1** |
| **A20** | **B-cell lymphoma** | **BALB/c** | **spontaneous** | **20** | **NR** |
| **CT26** | **colorectal** | **BALB/c** | **carcinogen** | **56** | **5** |
| **RENCA** | **renal** | **BALB/c** | **spontaneous** | **46** | **1.6** |
| **EMT6** | **breast** | **BALB/c** | **virus** | **13** | **1** |
| **EL4** | **T-cell lymphoma** | **C57BL/6** | **carcinogen** | **51** | **NR** |
| **MC38** | **colorectal** | **C57BL/6** | **carcinogen** | **75** | **5** |
| **LLCsr** | **lung** | **C57BL/6** | **spontaneous** | **72** | **5.3(LUAD)**<br>**7.5(LUSC)** |
| **B16F10** | **melanoma** | **C57BL/6** | **spontaneous** | **35** | **9.6** |
| **F9** | **teratocarcinoma** | **129S6/SvEv** | **embryoimplantation** | **11** | **NR** |

\*NR: not reported because there is no direct human equivalent data.

Mutational load has been correlated with tumor immune infiltrates (Rooney et al. 2015) and clinical response of checkpoint blockades in some human tumors (Snyder et al. 2014; Rizvi et al. 2015). We classified mutations into four categories using snpEff (Cingolani et al. 2012) based on their predicted impact on protein functions: HIGH (“The variant is assumed to have disruptive impact in the protein, probably causing protein truncation, loss of function or triggering nonsense mediated decay”), MODERATE (“A non-disruptive variant that might change protein effectiveness”), LOW (“Assumed to be mostly harmless or unlikely to change protein behavior”), or MODIFIER (“Usually non-coding variants or variants affecting non-coding genes, where predictions are difficult or there is no evidence of impact”). Next, we calculated mutational load for the HIGH and MODERATE mutations (Figure 1A), and compared with the nonsynonymous mutational load of the corresponding human tumors. In general, the mutational load for HIGH and MODERAT mutations in syngeneic models was higher than the median nonsynonymous mutational load in human tumors, although the values are within the range in human tumors (Table 1). MC38 has the highest mutational load, followed by LLCsr and CT26, while EMT6, F9 and 4T1 have the lowest mutational load. Carcinogen-induced models tend to have higher mutational burden, followed by spontaneously generated tumors, with viral induced models bearing the lowest mutational load (Figure 1A).

We focused on genes that lead to carcinogenesis when altered in human. We compared 43 point mutations across 27 genes whose human orthologs are reported in the TARGET database and annotated as actionable in OncoKB (http://oncokb.org/), as well as 8 variants of the tumor suppressor Trp53. Only four point mutations in two oncogenes (Kras G12C:LLCsr, G12D:CT26, G13D:EL4, and Nras Q61H:LLCsr) and four point mutations in two tumor suppressor genes (Pten T131P:B16F10, R130W:EL4, G209*:EMT6, and Trp53 E32*:LLCsr) matched in human tumors regardless of the tissue of origin (oncogene: exact variant; tumor suppressor genes: truncating mutation or missense mutations at the same amino acid) (Figure 1B). Next, we investigated whether genes frequently mutated in human tumors are also mutated in syngeneic models of the same tissue origin. While KRAS, APC and TP53 are frequently mutated in human colon tumors, CT26 only had homozygous Kras mutations (G12D, V8M), and MC38 only had Trp53 heterozygous mutations (G242V, S258I). In addition, SMAD4, mutated in approximately 12% of human colon cancer, was mutated in the MC38 model (Table 2). Neither colon syngeneic model contains an APC mutation, which is mutated in the majority of human colorectal cancer. Neither the breast-derived tumor model EMT6 nor 4T1 have activating mutations in PIK3CA: overall they contain fewer protein altering mutations than other syngeneic models, although 4T1 has an insertion in Trp53 that results in a frameshift mutation (E32fs). The LLCsr model also contains mutations in Trp53 (E32*, R334P) as well as Kras (G12C). Unlike CT26, the Kras (G12C) mutation in LLCsr is a heterozygous mutation. By contrast, the V600 BRAF mutation, a common mutation in human melanoma, was not identified in the melanoma B16F10 model. Similarly, genes frequently mutated in human kidney cancer such as VHL were not identified in the RENCA model.

**Table 2.** Frequently mutated human cancer genes and their mutations in syngeneic models of the same cancer type.

| Human Cancer | Human Gene(*) | Human Mutation Frequency | Model | Model Mutation | Model | Model Mutation |
| --- | --- | --- | --- | --- | --- | --- |
| BRCA | PIK3CA | 32.48 | 4T1 | NA | EMT6 | NA |
| BRCA | TP53 | 30.65 | 4T1 | p.Glu32fs | EMT6 | NA |
| BRCA | CDH1 | 11.41 | 4T1 | NA | EMT6 | NA |
| COAD | APC | 71.62 | CT26 | NA | MC38 | NA |
| COAD | TP53 | 53.6 | CT26 | NA | MC38 | p.Gly242Val<br>p.Ser258Ile |
| COAD | KRAS | 43.24 | CT26 | p.Gly12Asp<br>p.Val8Met | MC38 | NA |
| COAD | FBXW7 | 17.12 | CT26 | NA | MC38 | NA |
| COAD | PIK3CA | 14.86 | CT26 | NA | MC38 | NA |
| COAD | SMAD4 | 11.71 | CT26 | NA | MC38 | p.Gly351Arg |
| COAD | ATM | 11.26 | CT26 | NA | MC38 | NA |
| SKCM | BRAF | 51.23 | B16F10 | NA |  |  |
| SKCM | NRAS | 26.7 | B16F10 | NA |  |  |
| SKCM | ROS1 | 17.98 | B16F10 | NA |  |  |
| SKCM | ERBB4 | 16.35 | B16F10 | NA |  |  |
| SKCM | TP53 | 15.26 | B16F10 | p.Asn128Asp |  |  |
| SKCM | KDR | 13.35 | B16F10 | NA |  |  |
| SKCM | NF1 | 12.81 | B16F10 | NA |  |  |
| SKCM | CDKN2A | 12.26 | B16F10 | NA |  |  |
| LUSC | TP53 | 81.46 | LLCsr | p.Glu32*<br>p.Arg334Pro |  |  |
| LUSC | PIK3CA | 15.17 | LLCsr | NA |  |  |
| LUSC | CDKN2A | 14.04 | LLCsr | NA |  |  |
| LUSC | NF1 | 11.8 | LLCsr | NA |  |  |
| LUSC | ROS1 | 10.67 | LLCsr | NA |  |  |
| KIRC | VHL | 49.89 | RENCA | NA |  |  |

\*Genes from TARGET database with at least 10% mutation frequency in TCGA samples.

### Some syngeneic tumors displayed mesenchymal-like phenotype

In addition to genetic features, we compared the tumor histology of these mouse syngeneic models with human tumors. The *in vivo* tumors were stained with E-cadherin antibodies, an epithelial cell marker, and vimentin, a marker for cells undergoing epithelial to mesenchymal transition. Many models had high vimentin expression suggesting a more mesenchymal-like phenotype (Figure 2A, Supplemental Figure 3). In addition, the ratio of E-cadherin to vimentin is much lower than the corresponding human tumors in TCGA with the exception of RENCA (Figure 2B), suggesting that syngeneic models typically have a more mesenchymal-like tumor cellular phenotype than human tumors.

**Figure 3.**
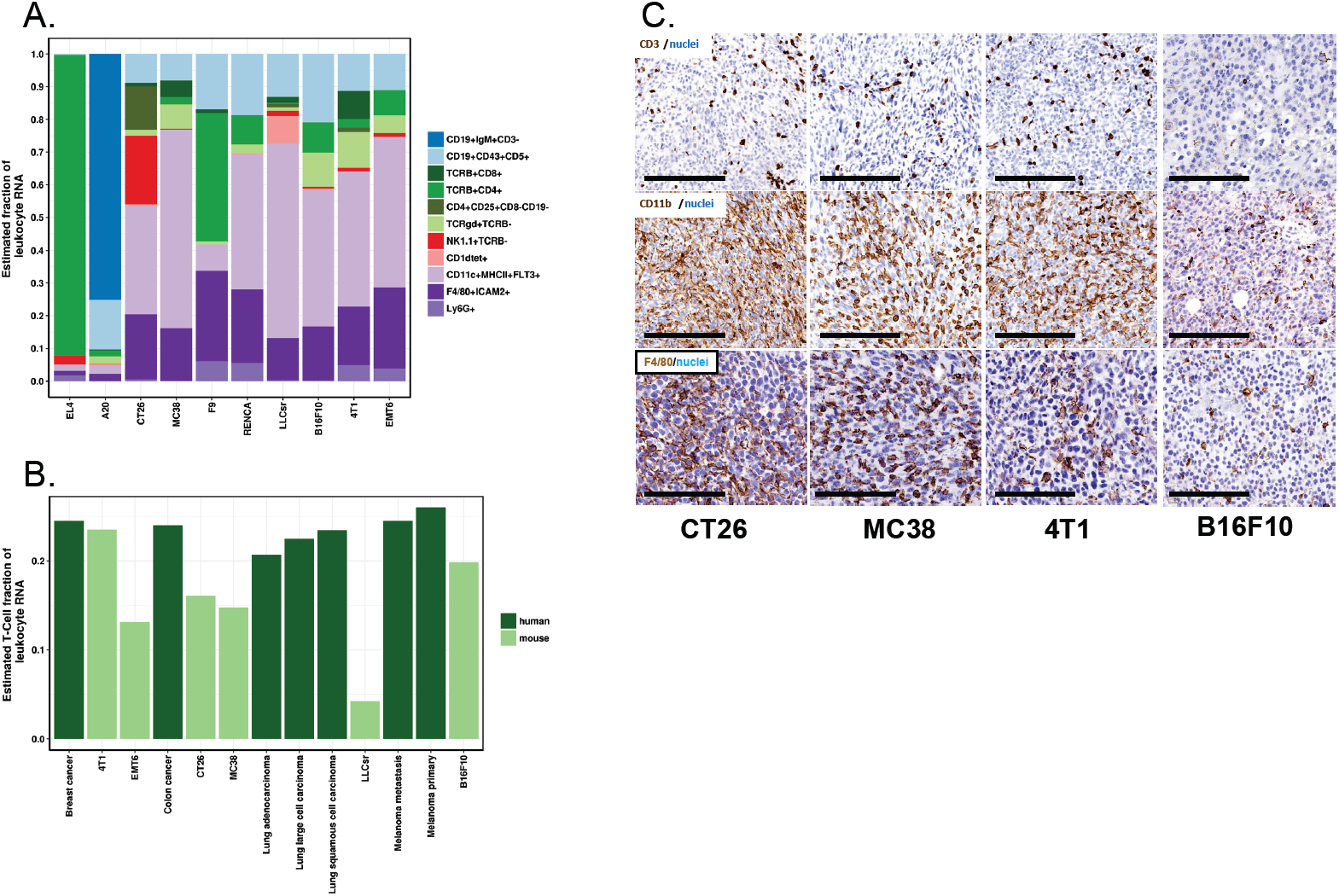
Immune subsets in syngeneic models. A. *In silico* immune cell deconvolution of syngeneic tumor samples. B. Comparison of estimated total T-cell fraction of leukocyte in selected mouse syngeneic models and their corresponding human tumors. Human data are downloaded from (Gentles et al. 2015). Total T-cell fraction plotted here is the sum of all predicted T-cell subsets including CD4+, CD8+, Treg, and gamma-delta T-cells. C. CD3 staining for T-cells, CD11b staining for myeloid cells, and F4/80 staining for macrophage.

### Syngeneic models have relatively low T-lymphocyte infiltration

The baseline immune infiltration of a panel of syngeneic models (Table 1) was evaluated by transcription profiling and chromogenic immunohistochemistry (IHC). We performed RNA-Seq for syngeneic tumors grown *in vitro* culture and *in vivo* (Supplemental Table S3), and implemented an *in silico* immune cell deconvolution using a nu-support vector regression (nuSVR) developed for mouse samples that is similar to approaches recently developed for human samples (Newman et al. 2015). As expected, a large percentage of T cells and B cells were predicted for EL4 and A20, T cell and B cell lymphoma models, respectively. A relatively high percentage of myeloid infiltration along with a relatively low percentage of T cells was predicted by *in silico* immune cell deconvolution (Figure 3A). The T-cell fraction was lower in most syngeneic models compared to the corresponding human tumors (Gentles et al. 2015) (Figure 3B). Furthermore, there were high levels of myeloid and macrophage infiltration by IHC in these models (anti-CD11b or anti-F4/80 staining, Figure 3C).

### Predicted neoantigen load in syngeneic mouse models does not correlate with cytolytic activity

Neoantigen load has been reported to correlate with tumor immune infiltrates (Rooney et al. 2015) and clinical response of checkpoint blockades in some human tumors (Snyder et al. 2014; Rizvi et al. 2015). We developed a neoantigen prediction pipeline based on MHC class I binding for the syngeneic models (details in method section); the number of predicted neoantigens correlates with mutational load (Supplemental Figure 4A), as in human tumors. Next, we evaluated the relationship between the predicted neoantigen load and tumor immunity using the cytolytic activity (CYT) as an indicator of the tumor immunity. We defined the cytolytic activity to be the log average (geometric mean) of two key cytolytic effectors, granzyme A (GZMA) and perforin (PRF1) (Rooney et al. 2015). Unlike what has been reported for human tumors, we did not observe a significant correlation between the neoantigen load and cytolytic activity (Supplemental Figure 4B).

**Figure 4.**
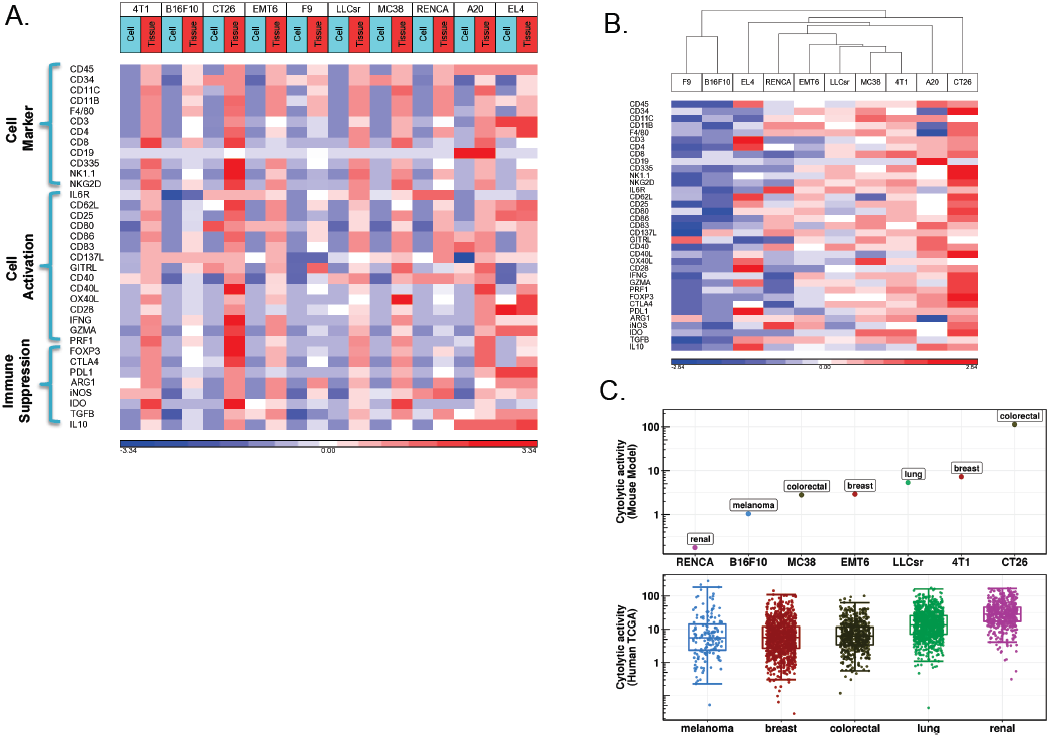
Immune infiltration in syngeneic models. A. Gene expression of immune cell type, immune cell activation and immune suppression markers in cells grown *in vitro* and tumor tissues from the transplantation. Gene expression shown as log2 of Transcript Per Million (TPM) and standardized across samples. B. Unsupervised clustering analysis of immune marker expression in tumor tissues from the transplantation separates syngeneic models into high and low infiltration models. C. Comparison of cytolytic activity of solid tumor syngeneic models with tissue matched human tumors from TCGA (human data were downloaded from (Rooney et al. 2015)). Cytolytic activity (CYT) is defined as the log-average (geometric mean) of Gzma and Prf1 expression in transcripts per million (TPM) as describe by Rooney et al

### Relative immunogenicity of syngeneic tumors display differences from that of the human tumors

We investigated the relative immunogenicity among syngeneic tumors using RNA-Seq and proteomics. Gene expression of many known markers of immune cell-type, immune cell activation and immune suppression were dramatically up-regulated in tumors *in vivo* compared to corresponding cells *in vitro*, consistent with immune infiltration (Figure 4A). In addition, unsupervised hierarchical clustering of gene expression of these immune-related genes displayed differential immune infiltration among models (Figure 4B). CT26, a colon cancer model, and 4T1, a breast cancer model, have the highest immune infiltration compared to other models while B16F10, a melanoma model and F9, a testicular teratoma, had lower infiltration. Likewise, total leukocyte infiltration in syngeneic models by CD45 (PTPRC) expression from RNA-Seq had a similar trend of immune infiltration. Cytolytic activity, another indicator of cancer immunity, was also highest in CT26 and 4T1 and lowest in B16F10 and RENCA among the solid tumor models (Figure 4C). CT26 was highly responsive to CTLA4 checkpoint inhibitors, but not to PD-1 inhibitors, while other models including the B16F10 melanoma model did not respond significantly to either of the checkpoint inhibitors (Supplemental Figure 5). The high immunogenicity of the CT26 model and low immunogenicity of B16F10 and RENCA models in our study differs from what has been reported in human tumors, where kidney cancer has the highest median cytolytic activity. Although human colon tumors and melanoma have similar median cytolytic activity, melanoma has a much more skewed distribution where a significant fraction of tumors have high cytolytic activity (Figure 4C). Our analysis suggested that the extremely high cytolytic activity of CT26 is primarily a result of dramatically higher expression of Gzma in this model (Supplemental Figure 6A). CT26 also had the highest cytolytic activity and Gzma expression based on our proteomic analysis, consistent with the RNA-Seq (Supplemental Figure 6B, Supplemental Figure 6C). The CT26 model was predicted to have significant NK cell infiltration based on *in silico* immune cell deconvolution of RNA-Seq (Figure 4A) which is consistent with high Gzma expression and corresponding high cytolytic activity, as Gzma has been previously shown to be expressed prominently in NK cells in mouse (http://www.immgen.org/).

**Figure 5.**
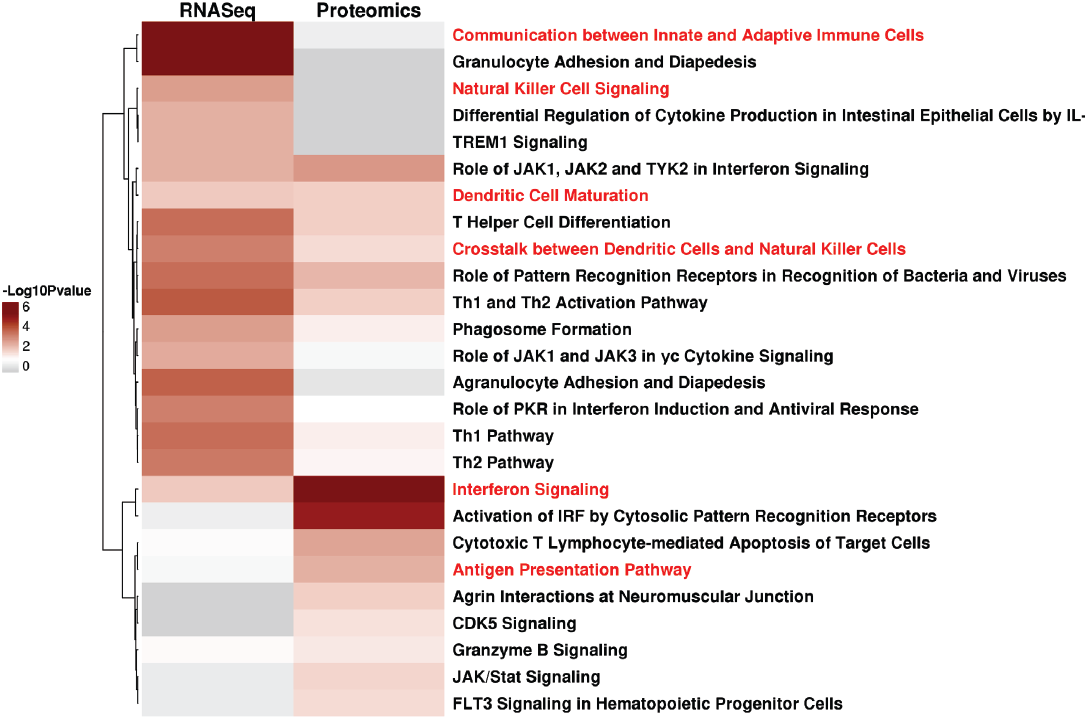
Top 10 significantly enriched pathways of genes up-regulated in CT26 *in vivo* tumor samples compared to *in vivo* tumor samples of other syngeneic models and CT26 *in vitro* samples from either RNA-Seq or proteomics data analysis (Fisher Exact p-value <= 0.05).

**Figure 6.**
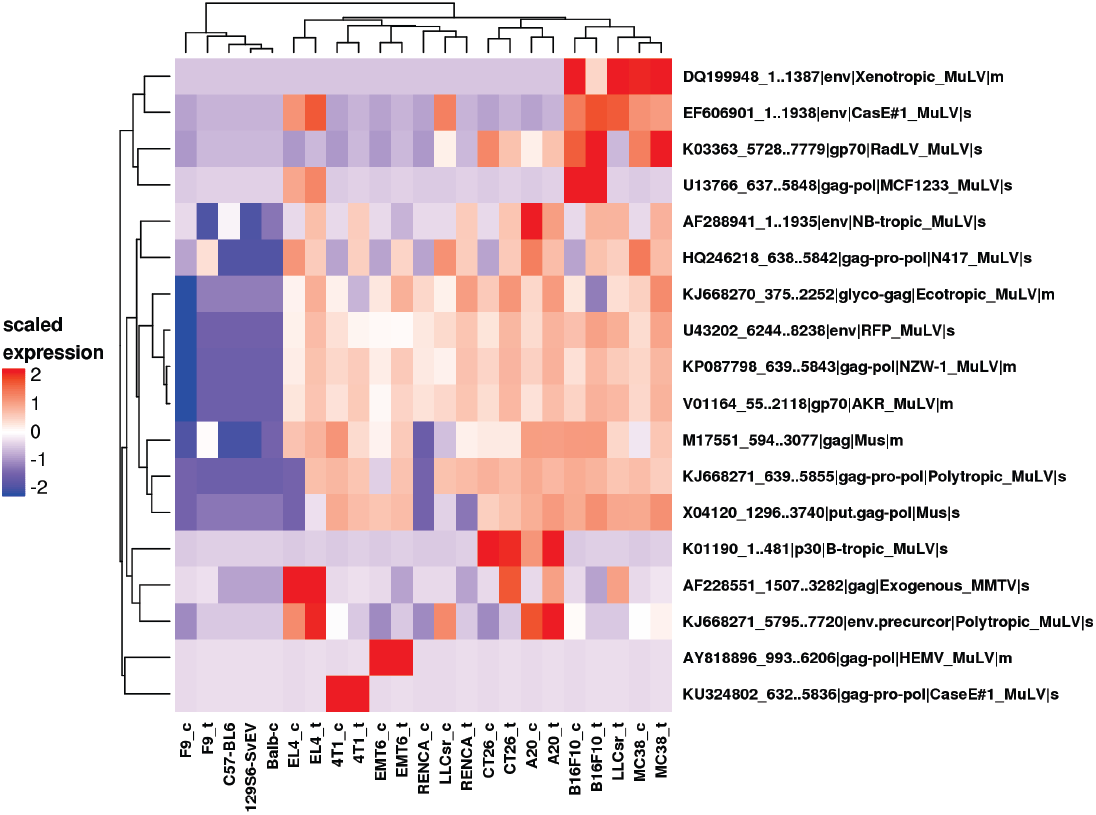
Viral peptides in syngeneic *in vitro* and *in vivo* samples from proteomic analysis (s: virus protein detected in soluble fraction; m: virus protein detected in membrane fraction, _t: *in vivo* tumor sample, _c: *in vitro* sample. Non-tumor samples are from tail of the parent strain). Hierarchical clustering using euclidean distance and complete linkage clustering method of log2 transformed and scaled LFQ values.

To further investigate the unique biology of the CT26 model, we analyzed pathways enriched in genes up-regulated in CT26 tumor samples *in vivo* compared to the same CT26 cells grown *in vitro* culture and other syngeneic *in vivo* tumors utilizing RNA-Seq: NK-related pathways including “Crosstalk between Dendritic Cells and Natural Killer Cells” and “Natural Killer Cell Signaling” were significantly enriched (Figure 5). Further analysis identified the “Crosstalk between dendritic cells and Natural Killer Cells”, “Interferon signaling”, and “Dendritic cell maturation” pathways as enriched both from RNA-Seq and proteomics. Our integrated pathway analysis is consistent with increased natural killer cell signaling in the CT26 model (Figure 5). Contrary to the large NK cell infiltration in the CT26 colon model, a much smaller fraction (approximately 1-3%) of NK cell infiltration has been reported in human colon tumors (Gentles et al. 2015).

### Proteomics characterization of virus antigen

Since viral antigens may also contribute to immunogenicity, we evaluated mouse viral proteins using custom proteomics. With the exception of LLCsr where a gag protein of mouse mammary tumor virus (ENA|AF228551_1507..3282) was detected in the tumor *in vivo* but not *in vitro* (Figure 6), 18 murine virus proteins were detected in cell lines when grown both *in vitro* and *in vivo*, with similar expression patterns across models as demonstrated by co-clustering of *in vitro* and *in vivo* samples for each model. 16 virus proteins were recurrent in more than one model while two (AY818896_993..6206, KU324802_632..5836) were expressed in only one model. One of the viral proteins that is expressed in 9 out of 10 models, murine leukemia virus envelope gp70 (ENA|V01164_55..2118), has been previously reported to be broadly expressed in murine cancer cells (Scrimieri et al. 2013). Interestingly, F9, a testicular teratoma, had very little virus protein expression compared to other models.

## Discussion

Syngeneic models are widely employed for evaluating efficacy, exploring mechanism of action, and generating predictive biomarker hypotheses to inform clinical development. While these models possess some molecular features similar to human cancers as assessed by next generation sequencing, proteomics and immunohistochemistry, they have many distinct properties: 1) only a small fraction of the somatic SNVs in mouse syngeneic models have the specific actionable mutation found in human tumors, 2) there is no significant correlation between neoantigen load and cytolytic activity in syngeneic models, 3) syngeneic models have lower T-lymphocyte infiltration compared to their corresponding human cancers, and 4) syngeneic models have a more mesenchymal-like phenotype than human tumors. Mouse models derived from a particular tissue typically do not reflect human tumors in the corresponding tissue. For example, the mouse colon tumor model CT26 has the highest immunogenicity and is the most responsive model to CTLA4 treatment among the models tested while the melanoma model B16F10 had the lowest immunogenicity and no response to the checkpoint inhibitors tested in our study, in contrast to the relatively high level of immunogenicity and response to checkpoint inhibitors for human melanomas. These differences suggest that mouse syngeneic models derived from a particular tissue do not represent features of human tumors from the same tissue; instead translation from mouse to human is more subtle and requires more in-depth analysis.

We investigated whether mouse syngeneic models are suitable for testing combination therapies spanning immune-oncology and genomically targeted agents by evaluating human cancer gene mutations in syngeneic models. First, many common mutations in human tumors are not represented in mouse syngeneic models (Table 2). Second, although some known human cancer genes are mutated in mouse syngeneic models, they rarely contain the same actionable variant observed in human tumors (Figure 1B). For example, of the three highly mutated genes, KRAS, APC and TP53 in human colon tumors, the CT26 colon tumor model only had activating Kras mutations (G12D, V8M), and the MC38 colon tumor model only had Trp53 mutations (G242V, S258I). While more than 70% of human colon tumors have mutations in APC, a common early event in the evolution of human colon cancer (Fearon 2011), neither of the two colon syngeneic mouse models contain APC mutations (Table 2). The lack of APC mutations in CT26 and MC38 suggest that these two colon syngeneic models are genetically distinct from human colon cancers. Similarly, the V600E BRAF mutation, a common oncogenic mutation in human melanoma, was not detected in the B16F10 melanoma model. These findings suggest that many of the syngeneic models may not fully recapitulate the genetic origin of human cancers, and these limitations may present challenges for studying combination therapies of immune-oncology and genomically targeted agents.

We analyzed the mutational and neoantigen load of ten syngeneic models by whole exome sequencing. While syngeneic models tend to bear a higher mutational load than the median mutational load observed in human tumors in TCGA, they are still within range (Table 1). Our analysis shows that mutational load is highly correlated with neoantigen load, suggesting that mutational load can be a surrogate for neoantigen load. However, unlike what has been reported in human, we did not observe a significant correlation of neoantigen load with immunogenicity or response to checkpoint blockade. This difference could be due to the difference in the activity of mouse surrogate antibodies relative to antibodies used in human studies, the limited number of models tested, or the diverse origin and tumor types represented in this study.

Beside the underlying genetic differences, we also observed cellular phenotypic differences between syngeneic mouse models and human tumors. First, we found that these models had a different morphological architecture compared to human tumors with more mesenchymal than epithelial phenotypes. Second, *in silico* deconvolution showed various immune infiltration patterns among these 10 models. The diversity of immune infiltration makes it possible to test the effect of therapies targeting different immune cells on the tumor growth. However, these models differ from human tumors with relatively low T-lymphocyte infiltration and high myeloid infiltration in several of the syngeneic models, along with a lower T-cell fraction than the corresponding human tumors.

In addition to neoantigens from somatic mutations, neoantigens from endogenous viral proteins can also act as tumor associated antigens and elicit CD8 T cell immunity. We identified viral proteins from two different types of oncogeneic viruses that are uniquely expressed in tumor models but not in skin epithelia from the tails of normal mice: murine leukemia virus and murine mammary tumor virus (Figure 6). Two of the viral proteins were selectively expressed in specific models, suggesting differences in pathogen exposure for each model when it was originally derived. While these murine specific viral antigens may play a role in cancer immunity in these models and lead to different responses to immunotherapy compared to human tumors, the specific immunity of these viral antigens remains to be investigated.

Our study suggested that the relative immunogenicity of various tumor types among syngeneic models differs from human tumors: the colon model CT26 is the most immunogenic while the melanoma model B16F10 is the least immunogenic. Recent studies (Spranger et al. 2015; Zou et al. 2016) implicated activation of the β-catenin oncogenic pathway as inducing resistance to anti-tumor immunity in melanoma. Consistent with this mechanism, the B16F10 model had high β-catenin expression while highly immunogenic models such as CT26 had lower expression (Supplemental Figure 7). Epigenetic silencing has also been reported to limit T cell immunity in ovarian and colon cancer (Peng et al. 2015; Nagarsheth et al. 2016; Zou et al. 2016). Ezh2, a member of the polycomb repressive complex 2 (PRC2) and Dnmt1, a DNA methylation enzyme were most highly expressed in the B16F10 model (Supplemental Figure 7), implicating both β-catenin pathway activation and epigenetic silencing as potential tumor intrinsic mechanisms leading to resistance to checkpoint inhibitors in this model. The B16F10 model may represent melanoma patients who do not respond to the checkpoint blockade, and may serve as a model for evaluating combination therapies of checkpoint inhibitors with agents that target these reported tumor intrinsic resistance mechanisms.

Unlike most human colon tumors, the CT26 colon tumor model had the highest immunogenicity among the models evaluated in our study. First, CT26 has high expression of immune markers and elevated cytolytic activity compared to other syngeneic models, consistent with previous reports of CT26 as a highly immunogenic model (Lechner et al. 2013). As noted previously, the high cytolytic activity in CT26 is largely attributed to high Gzma expression. In contrast to human, where Gzma can be expressed in both NK cells and CD8+ T cells, Gzma is predominantly expressed on NK cells in mouse (http://www.immgen.org/). Concordantly, our *in silico* immune cell type deconvolution identified significant NK cell infiltration in the CT26 model. Moreover, our integrated pathway analysis of both mRNA and protein expression identified several pathways related to NK cell function as highly enriched in CT26 tumors compared to either CT26 cells *in vitro* or other syngeneic tumors, providing further evidence of NK cell infiltration in the CT26 model that may contribute to its cytolytic activity. Further, antigen presentation and dendritic cell function are more active in CT26 compared to other models (Figure 5), and CD80 was expressed on CT26 cancer cells (Figure 4A). Besides its well-known function as a costimulatory molecule for T cell activation, CD80 has been reported to play a role in NK cell activation in both human and mouse cell lines expressing CD80 (Chambers et al. 1996; Luque et al. 2000). In addition, CD28 and CTLA4 expression has been reported in activated mouse NK cells, the interaction between CTLA4 and CD80 has a direct effect on IFN-γ release by NK cells, and CTLA4 expression has been reported in mouse tumor infiltrating NK cells (Stojanovic et al. 2014). Furthermore, we observed significant CTLA4 expression in CT26 *in vivo* tumor sample (Supplemental Figure 8) and on some tumor infiltrating NK cells based on single-cell RNA-Seq (data not shown). The role of NK cells in the remarkable response of CT26 to CTLA4 blockade (Supplemental Figure 8) as well as the potential mechanism of NK activation through CD80 expressed on CT26 cancer cells remains to be elucidated by future experiments.

In conclusion, we profiled the gene expression, proteomic, cellular phenotype, and pharmacology of several checkpoint inhibitors in ten syngeneic models. Our analysis suggested that these syngeneic models do not fully recapitulate human cancer biology, and the differences between the syngeneic models and human cancer may complicate translation of preclinical findings from these models to the clinic. Further model development to establish additional immune competent mouse models that harbor oncogenic driver mutations and encompass mutational loads more representative of human tumors may be required. While syngeneic models provide an opportunity to evaluate fundamental immunological pathways in the context of malignancy, they should be applied carefully with consideration of their differences from human tumors when informing clinical strategies.

## Methods

### Animals

Female inbred BALB/cAnNCrl 6-10 weeks of age were purchased from Charles River Laboratories (strain code 028). Female inbred C57BL/6J mice 6-10 weeks of age were purchased from Jackson Labs (strain 664). Female inbred 129S6/SvEvTac mice 6-10 weeks of age were purchased from Taconic Laboratories. All mouse strains were housed under specific pathogen-free conditions in Tecniplast IVC Green Line IVC cages in the vivarium of a Pfizer location in Pearl River, New York. Mice were housed on a 12:12 light:dark cycle, with *ad libitum* UV-sterilized water and low isoflavone 5V02 IF 50 irradiated Purina Chow (Purina). All animal studies were approved by the Pfizer Institutional Animal Care and Use Committee (IACUC) in accordance with the guidelines described in “Guide for the Care and Use of Laboratory Animals” (NRC, 2011).

### Syngeneic mouse models

4T1, A20, CT26, RENCA, EMT6, B16F10, and F9 cells were obtained from the American Type Culture Collection, Manassas, Virginia. MC38 cells were obtained from the laboratory of Antoni Ribas’ laboratory at UCLA, Los Angeles, California. LLCsr (Lewis lung carcinoma) cells were obtained from the laboratory of Shahin Rafii, Department of Genetic Medicine, Ansary Stem Cell Institute, Weill Cornell Medical College, New York, NY. For all models except the breast cancer model 4T1 and the colorectal model CT26, cells were injected in a 200 μl cell suspension in PBS in the right flank of 7-10 week old female mice. To establish the colorectal syngeneic model MC38, 1 x 10^6^ cells were implanted into C57BL/6J mice. To establish the lung cancer model LLCsr, the melanoma model B16F10, or the T cell lymphoma model EL4, 0.5 x 10^6^ cells were implanted into C57BL/6J mice. To establish the colorectal model CT26, 2 x 10^6^ cells in 50% Matrigel (Corning) were implanted into the right flank of 7-10 week old female BALB/cAnNCrl mice. To establish the breast cancer model EMT6, the B cell lymphoma model A20, or the renal cancer model RENCA, 1 x 10^6^ cells were implanted into BALB/cAnNCrl mice. To establish the 4T1 breast cancer model, 0.5 x 10^6^ cells were implanted subcutaneously into the right mammary fat pad of 7-10 week old female BALB/cAnNCrl mice. To establish the teratocarcinoma model F9, 2.0 x 10^6^ cells were implanted into 129S6/SvEvTac mice.

### Tumor collection for RNA-Seq and whole exome sequencing

When the calculated tumor volume was 400 −500 mm^3^, mice were euthanized using slow fill CO2 euthanasia according to Pfizer approved methods. The tumors were collected using aseptic technique and the tumors were transferred to RNAse and DNAse free tubes (Thermo Scientific Catalog 374320). Tumors were stored under liquid nitrogen until they were processed for RNA-Seq and WES.

### Whole exome sequencing

WES was conducted by Q^2^ solutions, USA using paired-end sequencing with read length of 2 x 100bps. Raw reads were aligned to the UCSC mm10 reference genome using BWA (v 0.7.5) (Li and Durbin 2009). Picard and GATK tools were used for duplicates removal, reads realignment and recalibration. Variants were called using both Varscan 2 (v2.3.6) (Koboldt et al. 2012) and SomaticSniper (v1.0.4) (Larson et al. 2012). Varscan 2 was performed using the *somatic* command with default parameters except --min-coverage was set to 20. The identified variants by varscan *somatic* were further filtered with *somaticFilter* and *processSomatic* commands using default parameters to obtain high confidence variant calls. Variant calls were made with SomaticSniper using default parameters and the results were further processed using scripts provided by the SomaticSniper package according to the suggestions from the manual to obtain high confidence variant calls. Variants from both Varscan 2 and SomaticSniper were annotated and filtered to obtain exonic variants using snpEFF (v4.1d)(Cingolani et al. 2012). Further, variants potentially leading to altered protein functions were defined as those annotated with MODERATE or HIGH IMPACT by snpEFF. The intersection of variant calls from VarScan 2 and SomaticSniper predictions was used as the final variant call list. 115 variants mapped to genes from TARGET database was further subjected to validation by Sanger sequencing at GeneWiz. Mutational load in exomes was calculated based on the identified HIGH and MODERATE impact mutations in protein-coding genes (assuming 32 Mb (Castle et al. 2014a) of protein-coding sequence). Median of Ts/Tv for each human tumor types was calculated based on data from Alexandrov et al. (Alexandrov et al. 2013). Median mutational load in exomes of human breast, lung squamous, lung adeno, colon, renal cell carcinomas and melanoma was calculated based on the identified nonsynonymous mutations in protein-coding genes (assuming 30 Mb (Alexandrov et al. 2013) of protein-coding sequence) using data downloaded from cBioPortal (http://www.cbioportal.org/). To evaluate the mutation of human known cancer genes in syngeneic models, genes from the TARGET (tumor <u>a</u>lterations relevant for <u>ge</u>nomics-driven therapy) database (https://software.broadinstitute.org/cancer/cga/target) were downloaded. Variants of cancer actionable genes were queried from OncoKB (http://oncokb.org/). Human mutation frequency data of TCGA samples for breast, lung squamous, colon, renal cell carcinomas and melanoma was downloaded from cBioPortal (http://www.cbioportal.org/). Subsequently, the mutation status of genes from TARGET database with at least 10% mutation frequency in TCGA samples was evaluated in syngeneic models that are of the same tissue origin.

### Neoantigen prediction

To predict neoantigens for each model, protein sequences for genes with predicted missense mutation were obtained from the Ensembl ftp site (ftp://ftp.ensembl.org/pub/release-84/fasta/mus_musculus/pep/). Two FASTA sequences were generated per variant site, wild type and mutant, with 10 amino acid sequences flanking each side of the variant site using pVAC-Seq (Hundal et al. 2016). The mouse haplotype (http://www.ebioscience.com/media/pdf/Mouse_Haplotype_Table.pdf) and candidate mutant epitopes for each variant were input to the IEDB MHC-I binding prediction tool. The IC50 for mutated epitopes with lengths of eight to eleven amino acids was predicted using NetMHCpan (Vita et al. 2015), and peptides predicted to have IC50 values less than or equal to 500 nM and more favorable than the wild type peptide were identified. We evaluated the expression of the corresponding gene for each predicted epitope and required that the gene expression to be above 2 TPM. Comparison of mutational load and predicted neoantigens was performed using the spearman method in R.

### Transcription profiling (RNA-Seq)

RNA-Seq profiling was conducted by Q^2^ solutions, USA. RNA from 30 cell cultures and 21 tumor tissue samples corresponding to 10 syngeneic mouse models were pair-end sequenced with read length of 2 x 100bps. Three replicates of cell culture and two replicates of tumor tissue were performed for each model. Raw reads were mapped to the UCSC mm10 reference genome using Bowtie 2 (v2.2.5) (Langmead and Salzberg 2012). Expected counts and normalized expression levels of genes in transcripts per million (TPM) were generated by RSEM (v1.2.20)(Li and Dewey 2011). Genes specifically up-regulated in CT26 *in vivo* tumor samples compared to CT26 grown in *in vitro* cell culture and *in vivo* tumor samples from other models were obtained as follows. First, genes significantly up-regulated in CT26 *in vivo* tumor samples compared to CT26 *in vitro* samples were identified using the DESeq2 package with criteria of adjusted p-value <= 0.01 and fold change >= 2. Secondly, genes up-regulated in CT26 *in vivo* tumors compared to *in vivo* tumors from other models were obtained using the DESeq2 package with criteria of adjusted p-value <= 0.01 and fold change >= 2. The final gene list was obtained from intersecting the above two gene lists and then was subjected to pathway analysis. All pathway enrichment analysis was performed using Ingenuity Pathway Analysis (IPA, Ingenuity® Pathway Analysis (IPA®)). To compare gene expression of markers of immune cell type, immune cell activation and immune suppression between cells grown *in vitro* and tumor tissues from the transplantation, standardized log2 (TPM) values were plotted in a heat map (Partek^®^ Genomics Suite^®^). Unsupervised hierarchical clustering analysis of gene expression of *in vivo* tumor samples was performed using the Partek^®^ Genomics Suite^®^ (Euclidean distance, average linkage clustering method). Cytolytic activity (CYT) was defined as the log-average (geometric mean) of Gzma and Prf1 expression value (TPM) as described by Rooney et al. (Rooney et al. 2015). Human cytolytic activity data were downloaded from Rooney et al. (Rooney et al. 2015). To compare ratio of the E-cadherin and vimentin gene expression between syngeneic models and human tumors, RNA-Seq data from TCGA were used. The ratio was calculated with the expression value (TPM) of E-cadherin and vimentin.

### *In silico* immune cell deconvolution

*In silico* immune cell deconvolution *of in vivo* tumor samples, either *in vitro* or *in vivo*, was performed on RNA-Seq profiling data using a nu-support vector regression (nuSVR) approach for mouse samples that is similar to approaches recently developed for human samples (Newman et al. 2015). To establish a murine immune cell-specific gene signature matrix, we downloaded RNA-Seq profiling data from 11 purified mouse immune cell subsets generated by the Immunological Genome Project (https://www.immgen.org/). These 11 immune cell subsets span all major hematopoietic lineages, and were double-sorted by flow cytometry from the spleen or peritoneal cavity of a 5-week old male C57BL/6J mouse (Jackson Laboratory). A list of the 11 immune cell subsets and the sorting markers are in Supplemental Table S4. More details about the RNA-Seq dataset can be found in the Sequence Read Archive (https://www.ncbi.nlm.nih.gov/sra) under accession PRJNA281360.

Raw RNA-Seq reads were aligned to the mouse reference transcriptome/genome (mm10) using Bowtie 2 (Langmead and Salzberg 2012) and summarized into gene-level transcripts per million (TPM) measures by RSEM (Li and Dewey 2011). TPM values were further quantile normalized before subsequent analysis. We extended a procedure for the selection and optimization of an immune cell specific gene signature matrix to mouse samples that is similar to approaches recently developed for human samples (Newman et al. 2015). Since there are no biological replicates per immune cell type, we used a Z-statistic to test whether any gene is significantly over-expressed in one immune cell subset versus all others. We kept all candidate genes for each cell type that have a q-value from the Z-test less than 0.01 and are expressed at least two-fold above the 3^rd^ quartile expression values of all genes in a cell type. An equal number of candidate genes from each cell type (sorted by expression fold change from the cell type of interest and the mean from all other cell types) were combined to form a gene signature matrix, where the optimal number of candidate genes was determined using a conditional number minimization procedure (Newman et al. 2015). This process selected a 577-gene signature matrix for the 11 mouse immune cell subsets.

With this immune cell-specific gene signature matrix, we performed deconvolution of bulk tumor profiles using a nuSVR algorithm (Schölkopf et al. 2000). Related methods for deconvolution of immune subset have been established for human samples (Newman et al. 2015). Our approach incorporates unique fingerprints derived for mice immune components. A deconvolution p-value was calculated for each sample which indicates whether there is significant presence of any immune cells (among the 11 immune cell subsets included in the gene signature matrix). A p-value cutoff of 0.1 was used to indicate significant deconvolution. *In vitro* tumor samples were used as negative controls for deconvolution as they should not have any immune cells, except for the EL4 and A20 hematological tumor models. Output from a significant deconvolution is relative fractions of the 11 immune cell subsets in the bulk tumor samples. The fraction of each immune cell subsets is relative to the total leukocyte content (e.g. CD45+ cells) in the sample and should sum up to 100%. Stacked bar charts were used to display these immune cell subset fractions as the average of biological replicates of a tumor model.

Total T-cell fraction is calculated as the sum of all predicted T-cell subsets (mouse: CD4+, CD8+, Treg, and gamma-delta T-cells; human: CD8+, CD4+ naïve, CD4+ memory RO unactivated, CD4+ memory RO activated, T cells follicular helper, T cells gamma delta, Tregs). Human leukocyte infiltration data are downloaded from (Gentles et al. 2015).

### Histology and Immunohistochemistry

Five micron sections were cut onto charged slides, dried, deparaffinized in xylene and rehydrated with graded alcohols to distilled H_2_O. For Hematoxylin and Eosin staining, sections were submerged in Tacha’s Auto Hematoxylin (Biocare Medical, Concord, CA, USA) for 1 min then rinsed in distilled H_2_O until clear. Slides were then submerged in tap water with agitation for 1 min followed by 1 min in 80% Reagent Alcohol (Thermo Fisher, Histoprep). Sections were then submerged in Eosin Y (Thermo Fisher) for 1.5 min followed by three 5 second washes in 95% Reagent Alcohol (Thermo Fisher, Histoprep), two 5 second washes in 100% Reagent Alcohol (Thermo Fisher, Histoprep), and finally in Xylene (Thermo Fisher, Histoprep) before being coverslipped with Permount mounting medium (Fisher Scientific Co. L.L.C., Pittsburgh, PA, USA). Immunohistochemistry heat-induced epitope retrieval was performed in the Retriever 2000 pressure cooker (Electron Microscopy Sciences, Hatfield, PA, USA) in Citrate Buffer pH 6.0 (Invitrogen, Carlsbad, CA, USA) or Borg buffer pH 9.5 (Biocare Medical, Concord, CA, USA) and cooled to room temperature for 20 min. Endogenous peroxidase activity was inactivated with Peroxidazed 1 (Biocare Medical, Concord, CA, USA) for 10 min. Non-specific protein interactions were blocked for 10 min with Background Punisher (Biocare Medical, Concord, CA, USA). Sections were incubated with primary antibodies, CD11b (AbCam, EPR1334,Citrate, 0.088μg/ml), F4/80 (Spring Bioscience, M4152, Citrate 2.5μg/ml), RA3-6B2, Borg, 2.5μg/ml), CD3 (Spring Bioscience, SP162, Borg, 1.33μg/ml), vimentin (Cell Marque, SP20, Citrate, 3μg/ml), E-cadherin (AbCam, EP700Y, Citrate, 0.18μg/ml), for 1 hr, washed in TBS and incubated with SignalStain Boost IHC Detection Reagent (Cell Signaling Technologies, Beverly, MA, USA) for 30 min. Following washes in TBS, immunoreactivity was visualized by development with 3,3’-diaminobenzidine (DAB+, Dako, Carpinteria, CA, USA) for 5 min. Immunostained sections were briefly counterstained with CAT Hematoxylin, washed in tap water, dehydrated in graded alcohols, cleared in xylene, and coverslipped with Permount mounting medium (Fisher Scientific Co. L.L.C., Pittsburgh, PA, USA).

### Proteomics acquisition and data analysis

All tissues were dissociated in 100 mM sodium carbonate using TissueLyzer II (QIAGEN) for 0.5 min at a rate of 0.30 repetitions three times and stored on ice for 30 sec between the dissociation cycles. The cells grown *in vitro* were detached with CellStripper (Mediatech), washed with cold PBS twice and cell pellets were collected. 100 mM sodium carbonate containing 1 mM DTT, pH 11.5 was used to lyse cells by incubating the cell suspension on ice for 60 min. For all tissue and cell protein lysates the pH was adjusted to pH 8 by the addition of Tris-HCl buffer at pH 7.0 and a final concentration of 2 mM MgCl2. Universal Nuclease (ThermoFisher) was used to dechromatinize nuclear DNA for 30 min on ice. The membrane fraction was isolated by centrifugation at 20,000 g for 60 min at 4°C and the supernatant was transferred to a fresh tube as the soluble fraction. The precipitated fraction following this centrifugation is membrane-protein enriched and is referred to as the “membrane” fraction, whereas supernatant is referred to as the “soluble” proteome fractions. The resulting membrane fraction (pellet) was solubilized using RIPA buffer (25 mM TrisHCl, pH7.6, 150 mM NaCl, 1% SDS, 1% sodium deoxycholate, 1% NP-40 and 1X protease inhibitor) to solubilize membrane proteins, whereas the soluble fraction was used directly for subsequent processing following determination of protein concentration by BCA assay. Each fraction was processed separately using filter-assisted sample proteolysis (FASP). Briefly, 50μg of protein was loaded on to a PES 100 KDa filter, washed 4 times with 8 M Urea in 100 mM Tris-HCl buffer, pH 8.5. Proteins were reduced with 5 mM DTT (freshly prepared) and alkylated with 10 mM idoacetamide in the dark. Samples were then washed three times with freshly prepared 25 mM ammonium bicarbonate buffer and digested with trypsin/LysC with protein:enzyme at a 25:1 weight ratio overnight. Following digestion, the peptides were captured by centrifugation at 15,000 g for 20 min, followed by adjusting the solution pH to acidic by adding 10% formic acid to the final concentration of 0.25% formic acid.

Each sample was acquired on a Thermo Scientific™ Q Exactive™ Hybrid Quadrupole-Orbitrap Mass Spectrometer fitted with a Dionex nano liquid chromatography and EASY-Spray™ Ion source. The tryptic digest was loaded onto a reversed-phase pre-column (C18 trap column, Acclaim PepMap 100, 100 μm x 2 cm, Thermo Fisher Scientific, Inc.). Peptide separation was conducted *via* nano-LC using a 75 μm x 150 mm PepMap C18 EASY-Spray column (3 μm, 100 Å particles, Thermo Scientific, Inc.). The gradient was comprised of an increase from 2 to 35% mobile phase B (0.1% formic acid in acetonitrile) over 160 min, followed by 35 to 80% B over 10 min and a hold at 80% B for the last 10 min, all at a fixed flow rate of 300 nl/min in an UltiMat 3000 RSLCnano system (Thermo Fisher Scientific, Inc.). Q Exactive runs were operated with data dependent top 10 method and the parameters were as follows: resolution 70,000 at m/z 200 for MS1 with a scan range of 300-1650 m/z, a predictive AGC target of 3 x 10^6^ and a maximum injection time of 100 ms; 17,500 at m/z 200 for dd-MS2 with a predictive AGC target of 2 x 10^5^, maximum injection time 120 ms, NCE of 25, 20s dynamic exclusion and underfill ratio 3%.

For soluble and membrane fractions, peptides were identified and quantified for label-free protein quantification from raw mass spectrometric files using MaxQuant software (version: 1.6.1.0) (Cox and Mann 2008), respectively. Database searching was performed in MaxQuant using the Andromeda search engine (Cox et al. 2011) against the mouse Uniprot database (57208 entries, May 2017 version) supplemented with our internal non-redundant virus protein sequences containing 903 ENA and 260 GenBank entries. Andromeda search parameters for protein identification were set as follows: maximum mass tolerance of 20 ppm for the first search, 4.5 ppm for the precursor ions after non-linear recalibration and 20 ppm for fragment ions; digestion was set to specific “Trypsin/P” allowing max missed cleavage events of two. Oxidation of methionine and protein N-terminal acetylation were set as variable modifications. Carboxyamidomethylation of cysteines was specified as a fixed modification. A total of three modifications were allowed per peptide. Minimal required peptide length was seven amino acids. MaxQuant LFQ “match between run” option was enabled with a match time window of 0.7 minutes after retention time alignment. “Requires MS/MS for label-free quantification (LFQ) comparisons” was not enabled to allow maximum MS peak features. Confidently identified proteins were required to have a minimum of two matched peptides with one peptide being uniquely matched at a false discovery rate (FDR) of less than 1%. Proteins were quantified by delayed normalization computed in MaxQuant’s label-free quantification option (Cox et al. 2014).

LFQ intensity data by protein group was further analyzed in Perseus (version: 1.6.0.7) (Tyanova et al. 2016). All proteins “identified by site”, “reverse” and contaminants were removed. LFQ intensity value was log2 transformed and the resulting data matrix was further filtered requiring minimal valid values of 100% in at least one model (Supplemental Table S5).

Proteins that are significantly over-expressed in CT26 *in vivo* tumor samples were obtained by pairwise comparison of the log2 transformed LFQ value of CT26 *in vivo* tumor samples from membrane fraction or soluble fraction to one of the other tumor models by applying a filter requiring a valid value of >=75% in at least one pair and using student t-test within Persues. Proteins with FDR (Benjamini-Hochberg) <= and fold change >= 2 in all comparisons as well as CT26 *in vivo* vs. CT26 *in vitro* samples were selected for further pathway and function enrichment analysis. If proteins were detected in both membrane and soluble fractions, the identification from the fraction with the higher LFQ values of the proteins will be kept. Pathway, function enrichment and network analysis were performed using Ingenuity Pathway Analysis (IPA, Ingenuity® Pathway Analysis (IPA®).

Hierarchical clustering of virus protein was performed within R using the Euclidean distance and the complete linkage clustering method. LFQ values of identified virus proteins were first log2 transformed and then scaled separately within *in vitro samples* and *in vivo* samples (which included both tumors and normal tails) before clustering analysis.

### Antibodies to mouse immune checkpoint proteins

Rat IgG2a to mouse PD-1/CD279 (clone RMP 1-14) was purchased from BioXcell and was dosed *in vivo* at 10 mpk i.v. q3dx3. Hamster IgG to mouse CTLA4/CD152 (clone 9H10) was purchased from BioXcell and was dosed *in vivo* at 10 mpk i.v. q3dx3.

### *In vivo* evaluation of antibodies to mouse immune checkpoint proteins

When the average tumor volume reached approximately 70 to 160 mm^3^, mice were randomized into treatment groups, with 10 mice in each treatment group. Antibodies or vehicle (PBS) were administered intravenously on study day 0 and then the animals were dosed once every 3 days for 3 doses. Tumors were measured 1 – 3 times per week and tumor volume was calculated as volume (mm^3^) = (width x width x length)/2.

## Acknowledgements

We gratefully acknowledge inputs from our colleagues James Hardwick, Timothy Fisher, Hui Wang, Manfred Kraus, Stephanie Shi, Jinwei Wang, Dave Looper, Nicole Streiner, Xiaorong Li and Lingqi Luo. The results published or shown here are in whole or part based upon data generated by the TCGA Research Network: http://cancergenome.nih.gov/.

---

**Supplemental Figure 1.**
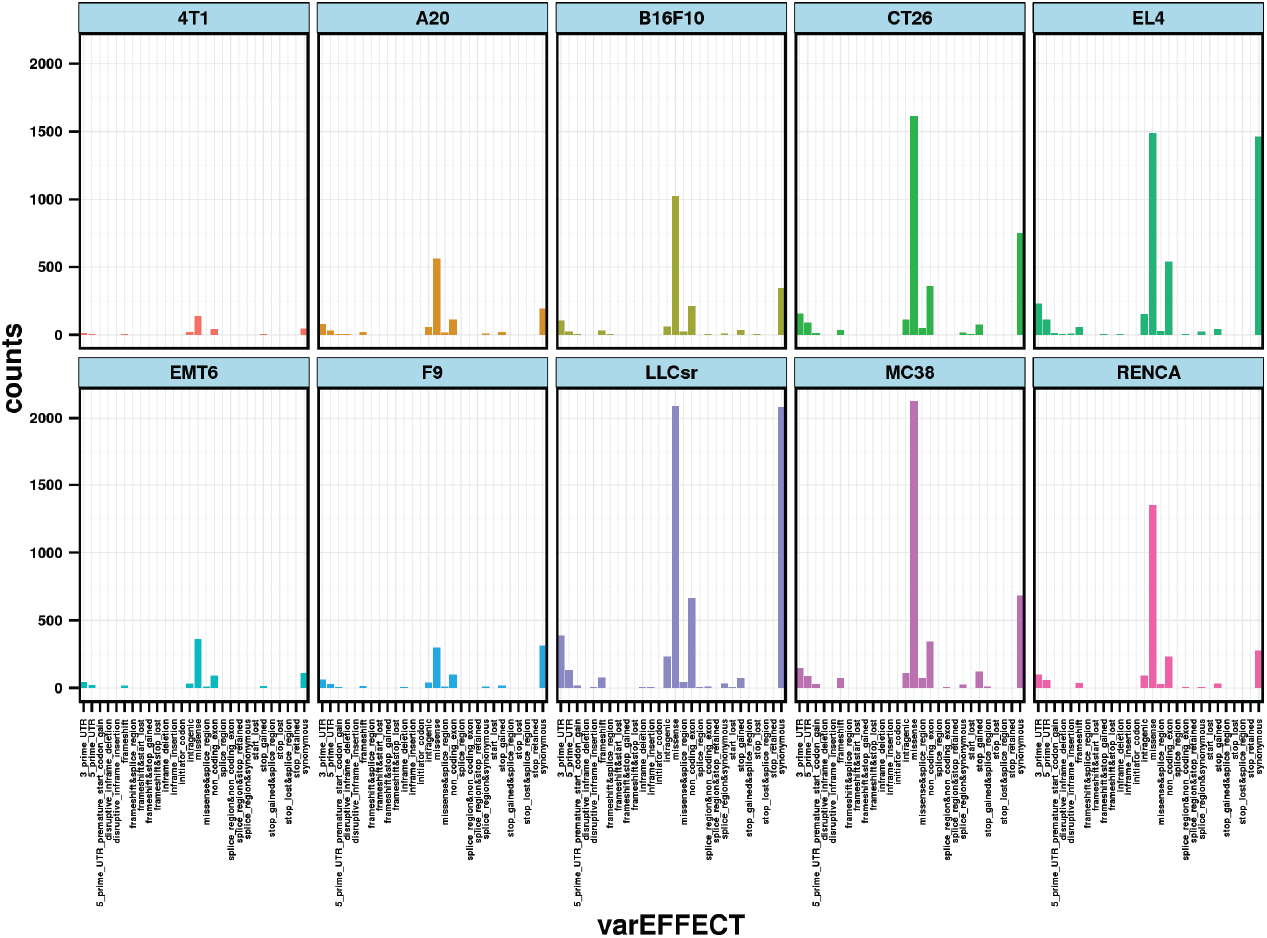
Mutation landscape of syngeneic models and types of somatic variants for each model.

**Supplemental Figure 2.**
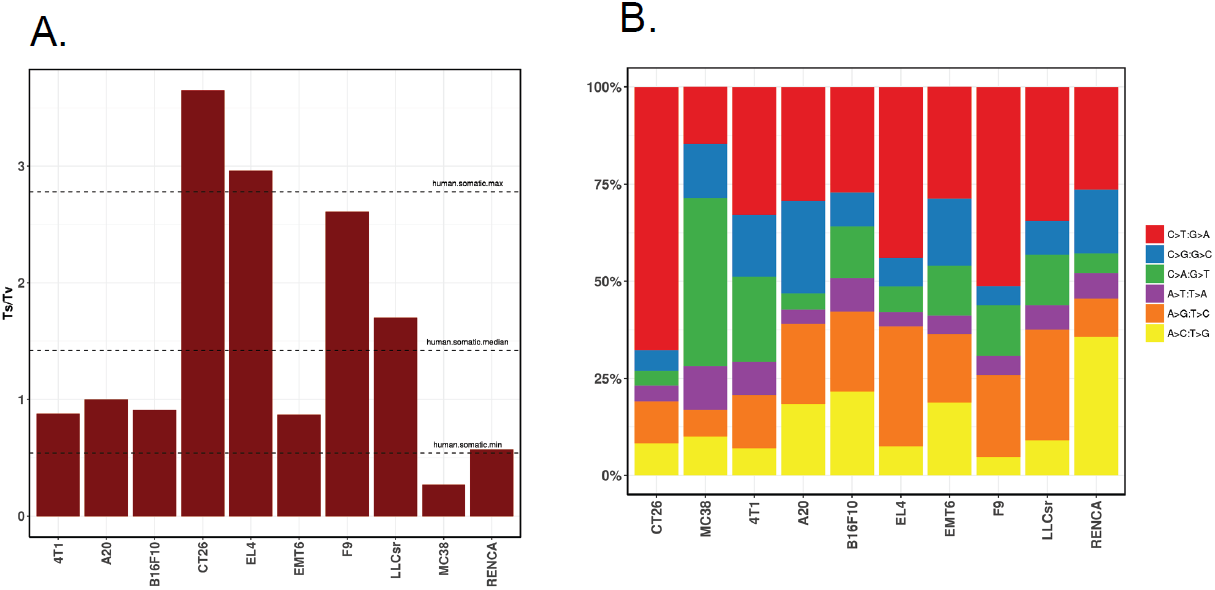
A. Ratio of Ts (Transition) to Tv (Transversion) substitution mutation in each syngeneic model; dashed lines represent maximum, median and minimum of median Ts/Tv from each human tumor types (data from (Alexandrov et al. 2013)) respectively. B. Single nucleotide mutation changes in each syngeneic model.

**Supplemental Figure 3.**
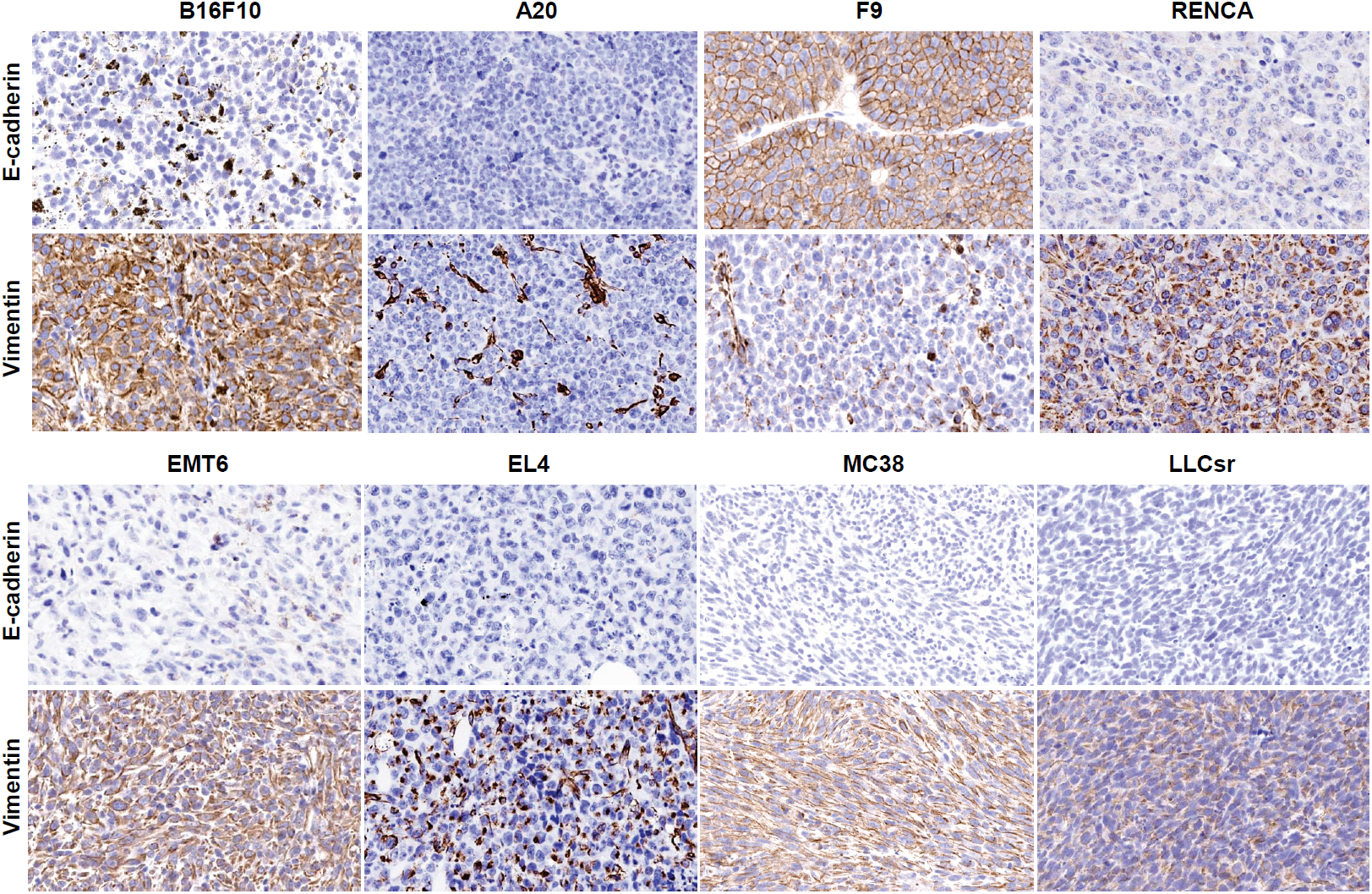
E-cadherin and vimentin stain in syngeneic models.

**Supplemental Figure 4.**
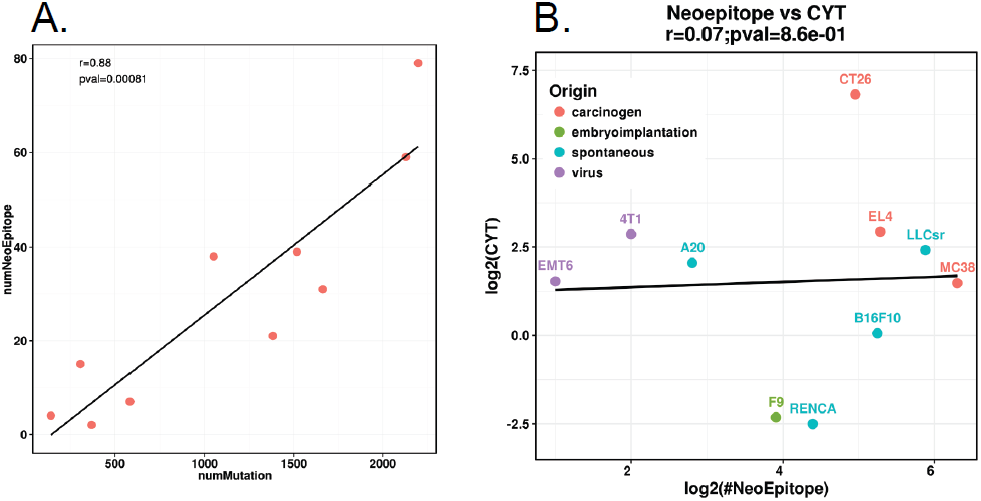
A. Correlation of neoantigen with mutational load in syngeneic models. B. Correlation of neoantigen with cytolytic activity in syngeneic models. The correlation was calculated using spearman method in R. numMutation: number of missense mutation.

**Supplemental Figure 5.**
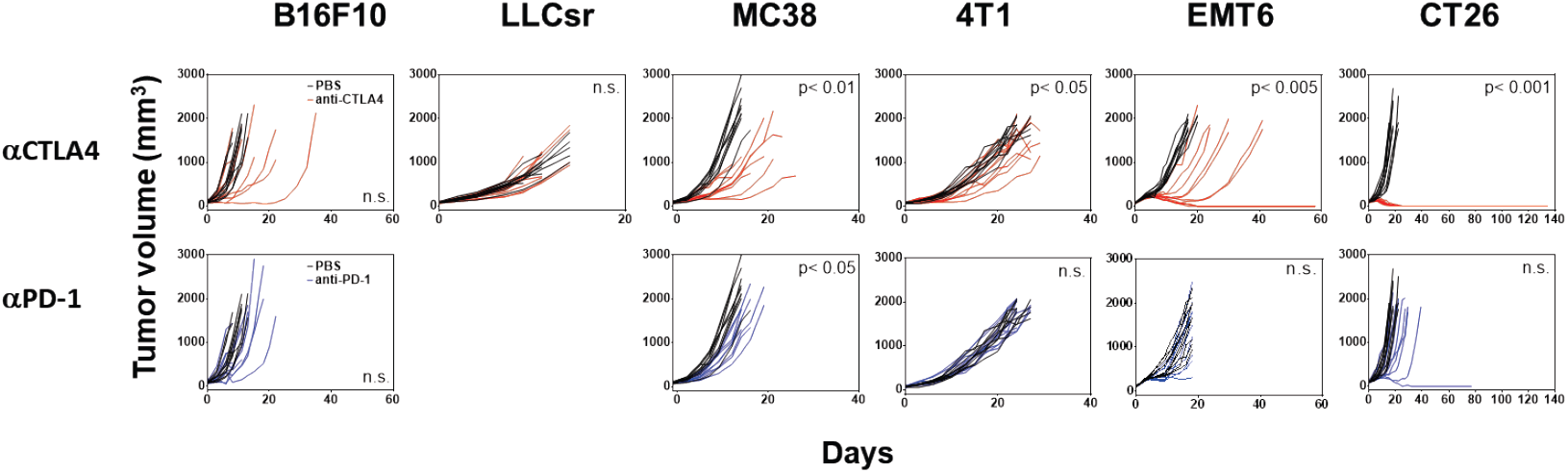
Response of syngeneic tumor models to anti-CTLA4, or anti PD-1. All mice were dosed intravenously. Individual tumor volumes are shown for 10 mice treated with PBS (black) or 10 mg/kg anti-CTLA4 (9H10) (red trace), or 10 mg/kg anti-PD-1 (RMP 1-14) (blue trace). All 10 mice bearing CT26 tumors dosed with anti-CTLA4 had no measurable tumor from study day 25 until measurements ended on study day 238. Comparison of mean tumor volumes was analyzed using log-transformed ANOVA. P-values are shown if there was a statistical difference between vehicle and antibody-treated groups. n.s. not significant.

**Supplemental Figure 6.**
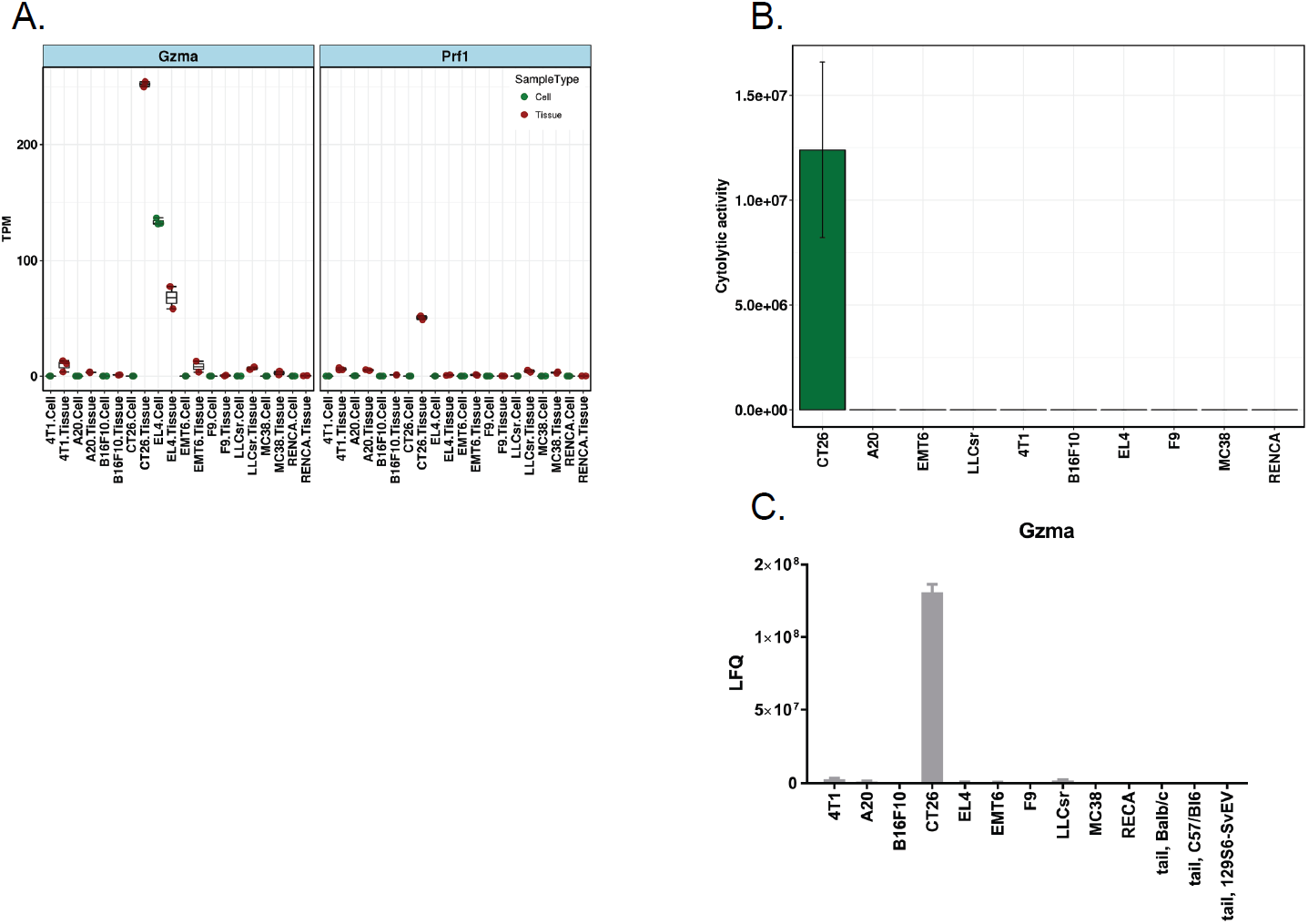
A. Expression of Gzma and Prf1 in syngeneic model grown *in vitro* (Cell) and *in vivo* (Tissue) across models based on protein expression. B. Cytolytic activity of syngeneic models. Cytolytic activity (CYT) is defined as the log-average (geometric mean) of Gzma and Prf1 protein expression. C. Protein expression of Gzma in soluble fraction. LFQ: label free quantitation.

**Supplemental Figure 7.**
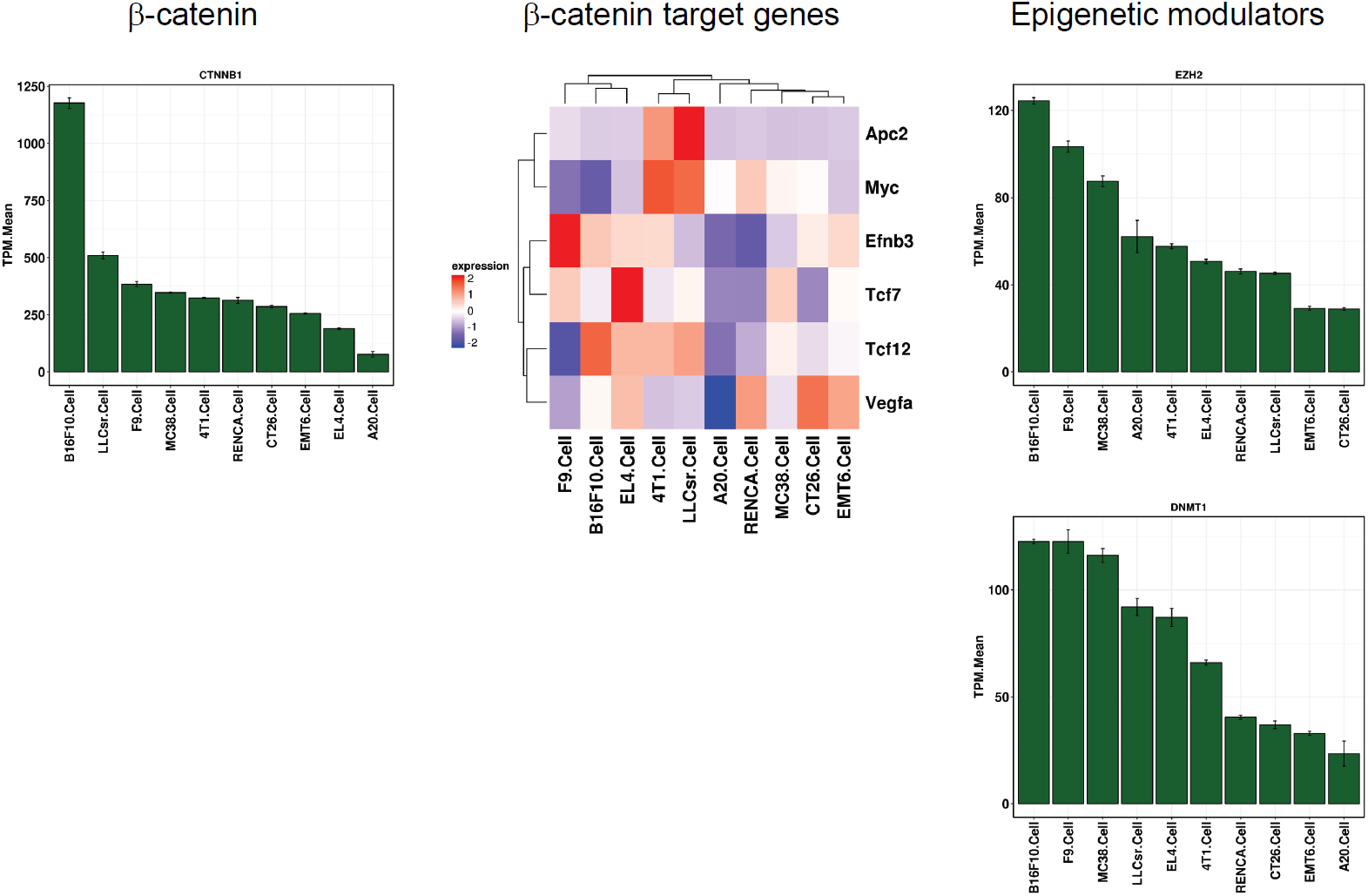
Gene expression of b-catenin, b-catenin target genes (gene list from (Spranger et al. 2015)) and epigenetic modulators (Ezh2, Dnmt1) across syngeneic models.

**Supplemental Figure 8.**
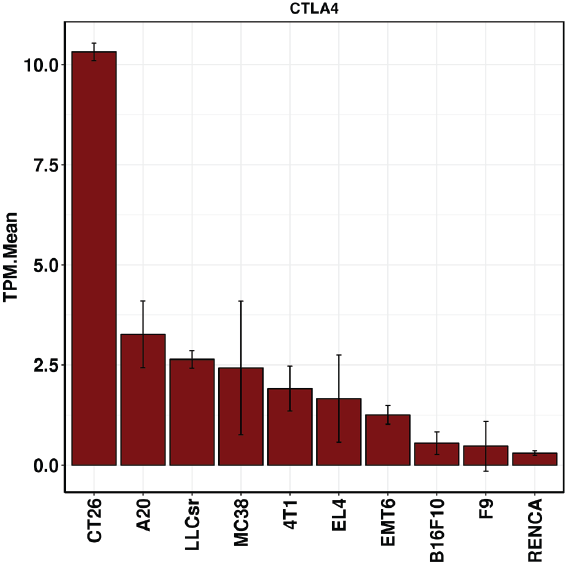
CTLA4 expression in syngeneic *in vivo* tumor samples.

## Reference

Alexandrov LB, Nik-Zainal S, Wedge DC, Aparicio SAJR, Behjati S, Biankin AV, Bignell GR, Bolli N, Borg A, Borresen-Dale A-L et al. 2013. Signatures of mutational processes in human cancer. Nature 500: 415–421.

Barretina J, Caponigro G, Stransky N, Venkatesan K, Margolin AA, Kim S, Wilson CJ, Lehar J, Kryukov GV, Sonkin D et al. 2012. The Cancer Cell Line Encyclopedia enables predictive modelling of anticancer drug sensitivity. Nature 483: 603–307.

Burns PA, Gordon AJE, Glickman BW. 1988. Mutational specificity of N -methyl-N -nitrosourea in the lacI gene of Escherichia coli. Carcinogenesis 9: 1607–1610.

Castle JC, Loewer M, Boegel S, de Graaf J, Bender C, Tadmor AD, Boisguerin V, Bukur T, Sorn P, Paret C et al. 2014a. Immunomic, genomic and transcriptomic characterization of CT26 colorectal carcinoma. BMC Genomics 15: 190.

Castle JC, Loewer M, Boegel S, Tadmor AD, Boisguerin V, de Graaf J, Paret C, Diken M, Kreiter S, Türeci Ö et al. 2014b. Mutated tumor alleles are expressed according to their DNA frequency. 4: 4743.

Chambers BJ, Salcedo M, Ljunggren H-G. 1996. Triggering of Natural Killer Cells by the Costimulatory Molecule CD80 (B7-1). Immunity 5: 311–317.

Cingolani P, Platts A, Wang LL, Coon M, Nguyen T, Wang L, Land SJ, Lu X, Ruden DM. 2012. A program for annotating and predicting the effects of single nucleotide polymorphisms, SnpEff. Fly 6: 80–92.

Cox J, Hein MY, Luber CA, Paron I, Nagaraj N, Mann M. 2014. Accurate Proteome-wide Label-free Quantification by Delayed Normalization and Maximal Peptide Ratio Extraction, Termed MaxLFQ. Molecular & Cellular Proteomics 13: 2513–2526.

Cox J, Mann M. 2008. MaxQuant enables high peptide identification rates, individualized p.p.b.-range mass accuracies and proteome-wide protein quantification. Nat Biotech 26: 1367–1372.

Cox J, Neuhauser N, Michalski A, Scheltema RA, Olsen JV, Mann M. 2011. Andromeda: A Peptide Search Engine Integrated into the MaxQuant Environment. Journal of Proteome Research 10: 1794–1805.

Dranoff G. 2012. Experimental mouse tumour models: what can be learnt about human cancer immunology? Nat Rev Immunol 12: 61–66.

Fearon ER. 2011. Molecular Genetics of Colorectal Cancer. Annual Review of Pathology: Mechanisms of Disease 6: 479–507.

Gao H, Korn JM, Ferretti S, Monahan JE, Wang Y, Singh M, Zhang C, Schnell C, Yang G, Zhang Y et al. 2015. High-throughput screening using patient-derived tumor xenografts to predict clinical trial drug response. Nature Medicine 21: 1318.

Gentles AJ, Newman AM, Liu CL, Bratman SV, Feng W, Kim D, Nair VS, Xu Y, Khuong A, Hoang CD et al. 2015. The prognostic landscape of genes and infiltrating immune cells across human cancers. Nature Medicine 21: 938.

Gould SE, Junttila MR, de Sauvage FJ. 2015. Translational value of mouse models in oncology drug development. Nat Med 21: 431–439.

Grosso JF, Jure-Kunkel MN. 2013. CTLA-4 blockade in tumor models: an overview of preclinical and translational research. Cancer Immunity 13: 5.

Gu Q, Zhang B, Sun H, Xu Q, Tan Y, Wang G, Luo Q, Xu W, Yang S, Li J et al. 2015. Genomic characterization of a large panel of patient-derived hepatocellular carcinoma xenograft tumor models for preclinical development. Oncotarget 6: 20160–20176.

Hundal J, Carreno BM, Petti AA, Linette GP, Griffith OL, Mardis ER, Griffith M. 2016. pVAC-Seq: A genome-guided in silico approach to identifying tumor neoantigens. Genome Medicine 8: 11.

Jenkins G, de G Mitchell I, Parry J. 1997. Enhanced restriction site mutation (RSM) analysis of 1,2-dimethylhydrazine induced mutations, using endogenous p53 intron sequences. Mutagenesis 12: 7.

Koboldt DC, Zhang Q, Larson DE, Shen D, McLellan MD, Lin L. 2012. VarScan 2: somatic mutation and copy number alteration discovery in cancer by exome sequencing. Genome Res 22.

Langmead B, Salzberg SL. 2012. Fast gapped-read alignment with Bowtie 2. Nat Meth 9: 357–359.

Larson DE, Harris CC, Chen K, Koboldt DC, Abbott TE, Dooling DJ. 2012. SomaticSniper: identification of somatic point mutations in whole genome sequencing data. Bioinformatics 28.

Lechner MG, Karimi SS, Barry-Holson K, Angell TE, Murphy KA, Church CH, Ohlfest JR, Hu P, Epstein AL. 2013. Immunogenicity of murine solid tumor models as a defining feature of in vivo behavior and response to immunotherapy. Journal of immunotherapy 36: 477–489.

Li B, Dewey CN. 2011. RSEM: accurate transcript quantification from RNA-Seq data with or without a reference genome. BMC Bioinformatics 12: 323–323.

Li H, Durbin R. 2009. Fast and accurate short read alignment with Burrows–Wheeler transform. Bioinformatics 25: 1754–1760.

Luque I, Reyburn H, Strominger JL. 2000. Expression of the CD80 and CD86 molecules enhances cytotoxicity by human natural killer cells. Human Immunology 61: 721–728.

Mosely SIS, Prime JE, Sainson RCA, Koopmann J-O, Wang DYQ, Greenawalt DM, Ahdesmaki MJ, Leyland R, Mullins S, Pacelli L et al. 2017. Rational Selection of Syngeneic Preclinical Tumor Models for Immunotherapeutic Drug Discovery. Cancer Immunology Research 5: 29–41.

Nagarsheth N, Peng D, Kryczek I, Wu K, Li W, Zhao E, Zhao L, Wei S, Frankel T, Vatan L et al. 2016. PRC2 Epigenetically Silences Th1-Type Chemokines to Suppress Effector T-Cell Trafficking in Colon Cancer. 76: 275-282.

Newman AM, Liu CL, Green MR, Gentles AJ, Feng W, Xu Y, Hoang CD, Diehn M, Alizadeh AA. 2015. Robust enumeration of cell subsets from tissue expression profiles. Nat Meth 12: 453–457.

Ostrand-Rosenberg S. 2004. Animal models of tumor immunity, immunotherapy and cancer vaccines. Current Opinion in Immunology 16: 143–150.

Peng D, Kryczek I, Nagarsheth N, Zhao L, Wei S, Wang W, Sun Y, Zhao E, Vatan L, Szeliga W et al. 2015. Epigenetic silencing of TH1-type chemokines shapes tumour immunity and immunotherapy. Nature 527: 249.

Rizvi NA, Hellmann MD, Snyder A, Kvistborg P, Makarov V, Havel JJ, Lee W, Yuan J, Wong P, Ho TS et al. 2015. Mutational landscape determines sensitivity to PD-1 blockade in non–small cell lung cancer. Science 348: 124–128.

Rooney MS, Shukla SA, Wu CJ, Getz G, Hacohen N. 2015. Molecular and genetic properties of tumors associated with local immune cytolytic activity. Cell 160: 48–61.

Schölkopf B, Smola AJ, Williamson RC, Bartlett PL. 2000. New Support Vector Algorithms Neural Comput 12: 1207–1245.

Sharma P, Allison James P. 2015. Immune Checkpoint Targeting in Cancer Therapy: Toward Combination Strategies with Curative Potential. Cell 161: 205–214.

Snyder A, Makarov V, Merghoub T, Yuan J, Zaretsky JM, Desrichard A, Walsh LA, Postow MA, Wong P, Ho TS et al. 2014. Genetic Basis for Clinical Response to CTLA-4 Blockade in Melanoma. New England Journal of Medicine 371: 2189–2199.

Spranger S, Bao R, Gajewski TF. 2015. Melanoma-intrinsic [bgr]-catenin signalling prevents anti-tumour immunity. Nature 523: 231–235.

Stojanovic A, Fiegler N, Brunner-Weinzierl M, Cerwenka A. 2014. CTLA-4 Is Expressed by Activated Mouse NK Cells and Inhibits NK Cell IFN-γ Production in Response to Mature Dendritic Cells. The Journal of Immunology 192: 4184–4191.

Tyanova S, Temu T, Sinitcyn P, Carlson A, Hein MY, Geiger T, Mann M, Cox J. 2016. The Perseus computational platform for comprehensive analysis of (prote)omics data. Nat Meth 13: 731–740.

Vizcaíno JA, Csordas A, del-Toro N, Dianes JA, Griss J, Lavidas I, Mayer G, Perez-Riverol Y, Reisinger F, Ternent T et al. 2016. 2016 update of the PRIDE database and its related tools. Nucleic Acids Research 44: D447-D456.

Yang Y, Yang HH, Hu Y, Watson PH, Liu H, Geiger TR, Anver MR, Haines DC, Martin P, Green JE et al. 2017. Immunocompetent mouse allograft models for development of therapies to target breast cancer metastasis. Oncotarget 8: 30621–30643.

Zou W, Wolchok JD, Chen L. 2016. PD-L1 (B7-H1) and PD-1 Pathway Blockade for Cancer Therapy: Mechanisms, Response Biomarkers and Combinations. Science translational medicine 8: 328rv324–328rv324.

